# Allelic expression imbalance of *PIK3CA* mutations is frequent in breast cancer and prognostically significant

**DOI:** 10.1101/2021.03.05.434137

**Authors:** Lizelle Correia, Joana M. Xavier, Ramiro Magno, Bernardo P. de Almeida, Filipa Esteves, Isabel Duarte, Matthew Eldridge, Chong Sun, Astrid Bosma, Lorenza Mittempergher, Ana Marreiros, Rene Bernards, Carlos Caldas, Suet-Feung Chin, Ana-Teresa Maia

## Abstract

PIK3CA mutations are the most common in breast cancer, particularly in the estrogen receptor positive cohort, but the benefit of PI3K inhibitors has had limited success compared with approaches targeting other less common mutations. We found allelic imbalances in the expression of *PIK3CA* in normal breast tissue and mapped a germline candidate regulatory variant. An imbalance was also frequently observed in the expression of the missense mutant and wild-type *PIK3CA* alleles in breast tumors from METABRIC and TCGA projects. Moreover, although 60% of tumors preferentially expressed the mutant allele, 10% did preferentially express the wild-type allele. Mechanistically, we show that these imbalances are more frequently due to regulatory variants in *cis* than altered copy-number and predict that somatic variants have a more significant role than germline ones. We further found that imbalanced allelic expression between mutant and wild-type alleles due to *cis-*regulatory variants associated with poor prognosis (*p*=0.0081).

Interestingly, ER^+^, PR^+^, and Her2^−^ tumors expressing very low levels of the mutant allele had the poorest prognosis (DSS <7.5yrs for ER^+^ and PR^+^ tumors and <5yrs for Her2^−^ tumors). Hence, our work provides compelling evidence to support the clinical utility of *PIK3CA* allelic expression in breast cancer in identifying this cohort of low mutant allele expressing patients of poorer prognosis, who will unlikely benefit from PI3K inhibitors. Furthermore, our work establishes a new model of differential regulation of critical cancer-promoting genes.

## Introduction

Activating oncogenic mutations are often characterized by gain-of-function single base alterations or focal DNA copy-number amplifications, where the gain of just a single copy of a mutant allele is sufficient for tumorigenesis [1]. These gains change the stoichiometric balance between mutant and wild-type alleles and are selected for in cancers, affecting approximately half of all oncogenic driver mutations [1]. Ultimately, they could dictate prognosis and therapeutic sensitivity.

However, the impact of gene dosage differences of oncogenic mutations generated at the gene expression level has been largely unexplored. Genetic variation and mutations regulate gene expression in an allele-specific manner – known as *cis*-regulatory variation [2] – by altering protein and miRNA binding. Normal *cis-*regulatory variation affects most of the human genome in all tissues and generates the wealth of phenotypic variation seen in a species [3–5]. Moreover, an extensive contribution from non-coding variants to RNA alterations was recently observed in tumors [6], including allelic imbalance of somatic mutations [7]. Nevertheless, one unsolved aspect is how much each mechanism contributes to generating allelic imbalances in expression and whether they do it independently or in synergy.

In breast tissue, germline regulatory variation affects frequently mutated genes [8] and is associated with disease risk. We and others have shown that variants affecting the expression levels of *BRCA1* and *BRCA2* modify the risk of breast cancer in germline mutation carriers [9,10]. We found that carriers of germline nonsense mutations in the tumor suppressor gene *BRCA2* were at a lower risk of developing breast cancer when the remaining wild-type allele was highly expressed.

Here, we hypothesize that *cis*-regulatory variation also modulates the penetrance of oncogenic coding mutations. In the context of a gene *cis*-regulated by a genetic variant generating imbalanced allelic expression, we postulate that an oncogenic activating mutation affecting the same gene will have a different clinical impact depending upon whether it occurs in the preferentially expressed allele or the less expressed one. We tested this model in the context of heterozygous mutations in *PIK3CA*, the most frequently mutated gene in breast cancer. Firstly, we investigated whether normal *cis*-regulatory variation regulated the expression of *PIK3CA* in normal breast tissue. Then, we calculated allelic expression ratios between mutant and wild-type copies in tumors from two large breast cancer datasets – METABRIC and TCGA – both normalized for DNA copy-number or not. Finally, we correlated the allelic expression ratios with clinical data. This approach allows us to distinguish between expression imbalances generated from *cis*-regulatory variation alone, altered DNA copy-number, or both mechanisms.

## Results

### Normal *cis*-regulatory variation affects *PIK3CA* expression in healthy breast tissue

To investigate whether *cis-*regulatory variation modulates the expression of *PIK3CA* in normal breast tissue (Figure 1A), we analyzed data from previous allelic expression analysis of normal breast tissue from 64 healthy donors [17]. We calculated the logarithm of the ratio of expression of one allele by the other in heterozygous SNP positions (aeSNPs) in all samples and defined differential allelic expression (DAE) when the difference between alleles was larger than or equal to 1.5-fold, i.e., the absolute value of log_2_ allelic expression ratio ≥ 0.58 (See Methods). A significant advantage of this approach is its robustness to detect *cis-*acting variant effects, as it cancels out the *trans* effects that might be acting on the same gene and influence both alleles equally. We found six SNPs in *PIK3CA* displaying DAE (daeSNPs) out of 14 aeSNPs tested (Figure 1B). All daeSNPs, except rs3729679, are in strong linkage disequilibrium (LD) with each other (Supplementary Table 1). rs3960984 showed the largest proportion of heterozygotes displaying DAE (57%), while three daeSNPs shared the smallest fraction (14%): rs12488074, rs4855093, and rs9838411.

**Figure 1.**
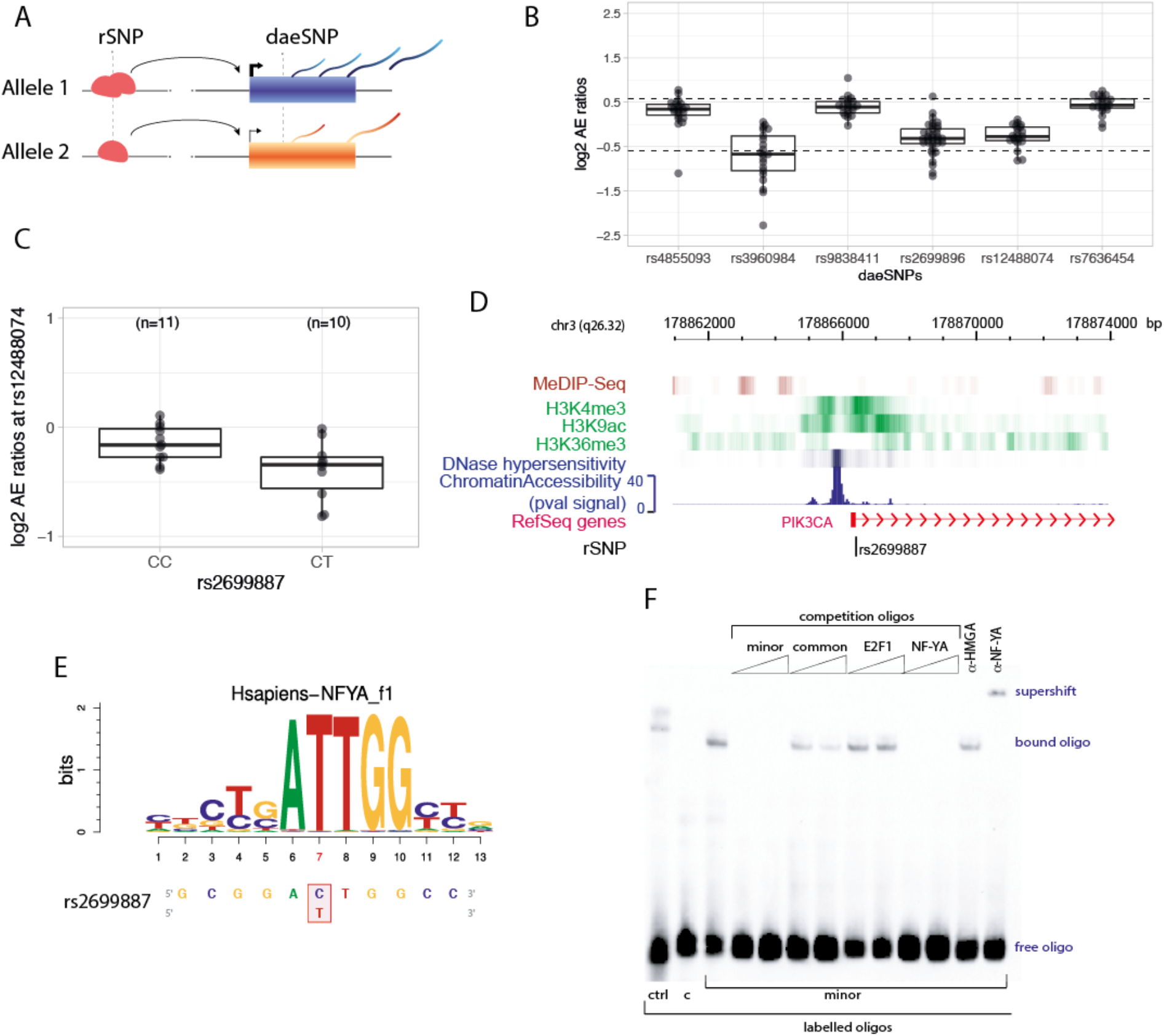
*Cis-*regulatory variation impacts on *PIK3CA* gene expression in normal breast tissue. **A** – Schematic representation of *cis-*regulatory SNP (rSNP) generating DAE on a target gene. Different alleles of an rSNP may, for example, have differential binding affinities to transcription factors (red), causing differences in the expression of both alleles detectable in heterozygous individuals for the daeSNP. **B** - DAE ratios for six daeSNPs in the *PIK3CA* gene region, each dot is a heterozygous individual for the corresponding variant indicated in the x-axis, dotted lines delimit the levels of 1.5-fold difference for either allele preferential expression (|DAE|= 0.58). **C** – rs2699887 significantly associates with DAE for rs12488074. Indicated p-values correspond to Student’s *t*-test test, with 1000 permutations. **D** – Genomic view of the location of rs2699887, showing methylation of surrounding CpG sites (MeDIP-seq), typical active promoter chromatin modification marks (H3K4me3, H3K9ac, H3K36me3), and open chromatin status (*DNAse*I hypersensitivity and Chromatin Accessibility p-val). Data from the Roadmap of Epigenomics project for breast HMECs. **E** – PWM analysis suggested rs2699887 disrupts an NF-YA binding motif at the four-nucleotide core sequence. **F** – rs2699887 differentially binds transcription factor NF-YA in vitro. Representative EMSA analysis using biotin-labeled oligonucleotides containing either rs2699887-C or rs2699887-T alleles, and one positive control (ctrl), using protein extracts of breast cancer cell line HCC1954. Competition assays included one (100x) or two concentrations (10x, 100X, as indicated by the gradient symbol) of unlabeled oligonucleotides, which included consensus binding sequences for NF-YA and E2F1. Supershift assays were carried out using monoclonal antibodies against NF-YA and one negative control (HGMA).

The range of allelic expression imbalances detected was between 1.55 and 2.14-fold (DAE ratio between 0.63 and 1.10). In four out of the six daeSNPs (rs7636454, rs3960984, rs12488074, rs9838411), the DAE ratios showed a unilateral distribution, with all samples with DAE preferentially expressing the same allele. For the other two daeSNPs, only one sample for each differed in the preferentially expressed allele. These patterns of DAE distribution suggest that the daeSNPs and the possible functional regulatory SNPs (rSNPs) are in strong, yet incomplete, LD with each other [18].

### Candidate rSNP for *PIK3CA* affects an NF-YA binding site at its promoter

Next, we identified the rSNPs acting on *PIK3CA* by carrying mapping analysis for all daeSNPs, using genotyping information from the SNPs located within a 250kb window from the gene limits (NM_006218, GRCh38/hg38). Initially, we identified a list of 44 candidate SNPs that, together with their 229 proxy SNPs, significantly explained the DAE observed (See Methods, Supplementary Table 2).

Functional analysis was carried for these 273 candidates, starting with *in-silico* analysis to identify those with greater *cis-*regulatory potential, followed by *in-vitro* validation. One strong candidate rSNP, rs2699887, was significantly associated with DAE detected at rs12488074 (permuted *p =* 0.03) (Figure 1C) and is an eQTL (expression quantitative trait locus) for *PIK3CA* (*p =* 0.012, Supplementary Figure 1) in tumors from METABRIC^24^, with the T minor allele associated with higher expression. rs2699887 resides in the first intron of *PIK3CA*, in a region rich in epigenetic marks and classified as an active promoter in breast mammary epithelial cells (vHMEC) and breast myoepithelial primary cells (Figure 1D). The motif position weight matrix (PWM) analysis ^19,20^ of the sequence surrounding rs2699887 suggested that NF-Y proteins could have preferential binding affinity to the minor allele (T allele) (Figure 1E). Electrophoretic mobility shift assays (EMSA) confirmed the preferential binding potential of the transcription factor NF-YA to the minor allele of rs2699887 (T allele) (Figure 1F).

### Preferential expression of the *PIK3CA* mutated alleles is frequent in breast tumors

Changes in DNA copy number in tumors are associated with changes in gene expression in *cis* [1,11–14], leading to dosage imbalances of coding mutations [7]. However, these differences can also be due to germline and somatic *cis-*regulatory variation, but their effect on mutation dosage imbalance is unknown. So, we set out to assess whether *PIK3CA* somatic mutations would have their functional effects modified by imbalances in allelic expression generated by regulatory variants, including the one we identified in normal breast tissue. We hypothesized that preferential expression of a gain-of-function mutation would have a more substantial clinical impact than those occurring in lowly expressed alleles, thus generating inter-tumor clinical heterogeneity (Figure 2A). To test this, we carried mutant vs. wild-type allelic expression analysis in breast tumor samples carrying somatic *PIK3CA* missense mutations on two independent sets of data, the METABRIC (n=94) and the TCGA (n=178) projects. Supplementary Table 3 presents a summary description of the two sets.

**Figure 2:**
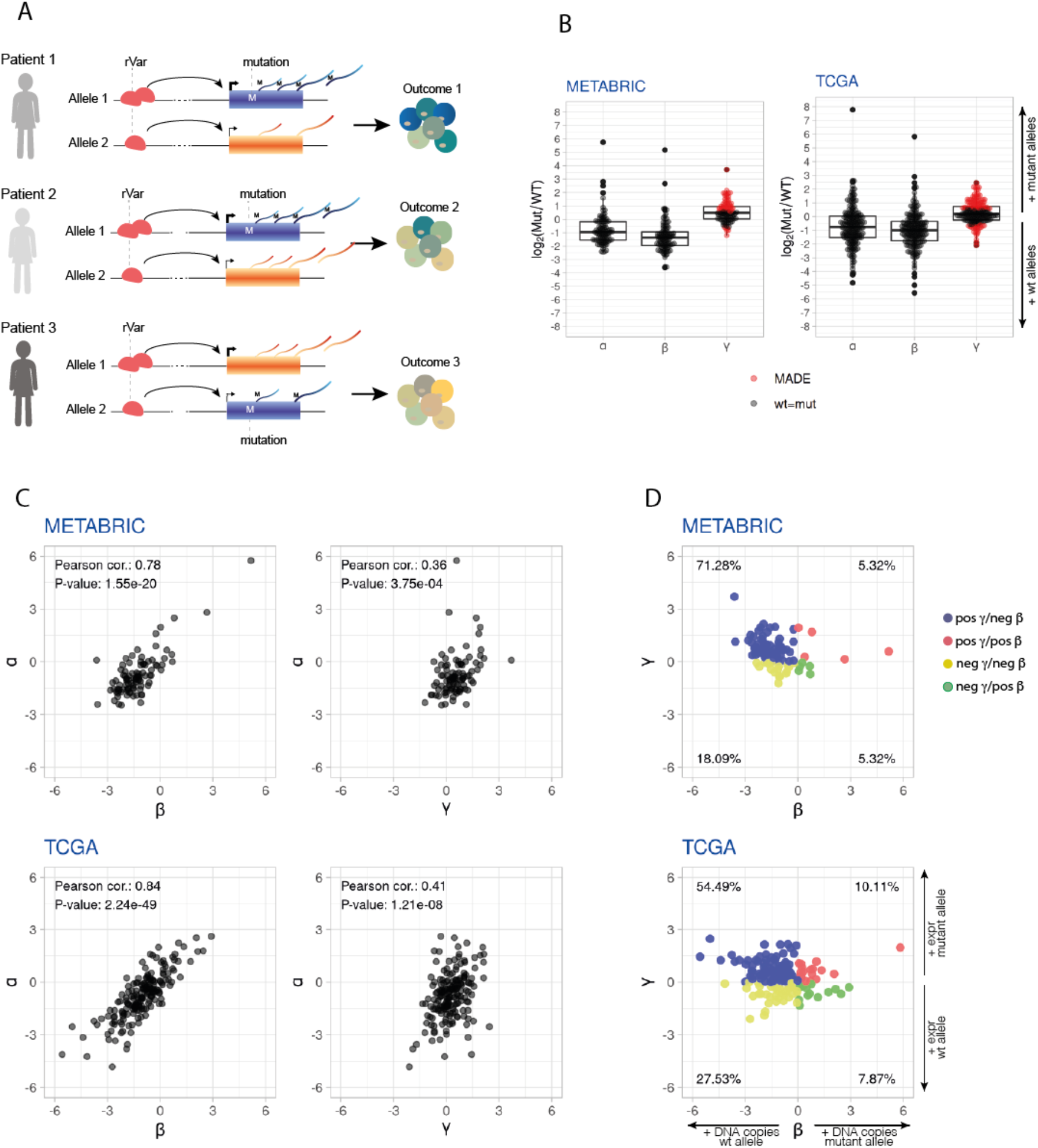
Mutant allelic imbalance in gene expression of somatic missense *PIK3CA* mutations is frequent in breast tumors, particularly for preferential expression of the mutant allele. **A** – Schematic representation of the hypothesis: *cis*-acting regulatory variants (rVar), either from germline or somatically acquired, generate different relative allelic expression ratios of mutant and wild-type alleles, resulting in tumors of different prognosis**. B** - Distribution of α, β and γ ratios in breast tumors. Tumors with MADE, |γ| ≥ 0.58, are displayed in red. **C** – Correlation analysis of α vs. β and α vs. γ, showing that both genomic copy-number dosage and allelic expression regulation contribute to allelic imbalances in the expression of mutations in tumors. **D** – Comparison of matched β and γ values, showing predominance of tumors with a preferential allelic expression of the mutated allele, albeit a higher number of wild-type allele copies.

Next, we calculated three allelic ratios from DNA-seq and RNA-seq data for each mutation:

α = log_2_ (number of mutant RNA-seq reads / number of wild-type RNA-seq reads), i.e., the net mutant allele expression imbalance;

β = log_2_ (number of mutant DNA-seq reads / number of wild-type DNA-seq reads), i.e., the mutant allele relative copy-number;

γ = α - β, i.e., the net mutant allele expression imbalance normalized for the DNA allelic copy-number imbalances, which corresponds to a putative mutant allele expression imbalance due to *cis-*regulation.

In this way, α reports on the overall allelic expression imbalance, generated by different mechanisms including copy-number aberrations, cellularity differences, and *cis-*regulatory variation, while γ reports specifically on the contribution from *cis-*regulatory variation (rVar in Figure 2A), including normal genetic variation, somatic non-coding mutations, and allelic epigenetic changes. Figure 2B displays the distributions of the different ratios.

We found that overall mutant allele expression imbalances (α ratio) are frequent in breast tumors, as seen for METABRIC (75.5%) and TCGA (68%) (Table 2). The same is true for γ ratios, at 50.4% for METABRIC and 46% for TCGA, indicating that *cis-*regulatory effects acting on mutations are also frequent in breast tumors. In both sets, we found samples with striking overall preferential allelic expression for the mutant allele (maximum 54-fold and 220-fold in METABRIC and TCGA, respectively), but not so for the preferential expression of the wild-type allele (fold differences of 5.5 and 28.2 in METABRIC and TCGA, respectively) (Table 2, Figure 2B). Similarly, the mutant allele’s more pronounced preferential expression trend was found for the γ ratio, 13 and 5.5-fold for METABRIC and TCGA, respectively, albeit with smaller fold differences between alleles.

Interestingly, we observed that within the samples with mutant allele differential expression (which we denominated MADE, defined as |γ| ≥ 0.58) there was a significant prevalence of samples that preferentially expressed the mutated allele in both data sets (binomial test Prob = 0.88, CI95% = [0.75 - 0.95], *p* = 1.0e-07 for METABRIC and Prob = 0.70, CI95% = [0.58 - 0.80], *p* = 7.6e-04 for TCGA).

**Table 1.** DAE of normal breast samples in six daeSNPs located at the PIK3CA region.

| daeSNP | Heterozygotes with DAE * | Preferentially Expressed Allele | Mean DAE * | SD | Min-Max | *P* t.test (one sample) |
| --- | --- | --- | --- | --- | --- | --- |
| rs7636454 | 24% (5/21) | A | 0.68 | 0.04 | 0.63-0.76 | 5.70E-06 |
| rs3960984 | 57% (12/21) | G | 1.11 | 0.46 | 0.64-2.28 | 5.91E-06 |
| rs3729679 | 20% (7/35) | G | 0.85 | 0.21 | 0.61-1.17 | 6.07E-05 |
|  | 3% (1/35) | A | 0.63 | 0 | 0.63-0.63 | - |
| rs12488074 | 14% (3/21) | G | 0.74 | 0.09 | 0.61-0.82 | 8.06E-03 |
| rs4855093 | 9% (2/22) | A | 0.74 | 0.04 | 0.70-0.78 | 3.23E-02 |
|  | 5% (1/22) | G | 1.1 | 0 | 1.10-1.10 | - |
| rs9838411 | 14% (3/22) | A | 0.76 | 0.2 | 0.59-1.04 | 3.34E-02 |

\* DAE calculated as $\log_2(\text{expression of allele 1}/\text{expression of allele 2})$ ; SD – Standard Deviation; *P* - p-value

**Table 2.** Summary of allelic expression imbalance of somatic *PIK3CA* mutations in breast tumors.

| Variable | N | METABRIC, N = 94<sup>†</sup> | TCGA, N = 178<sup>†</sup> |
| --- | --- | --- | --- |
| **alpha** | 272 | -0.96 (-2.47, 5.75) | -0.76 (-4.82, 7.78) |
| **alpha_group** | 272 |  |  |
| > wt |  | 63 (67%) | 96 (54%) |
| neutral |  | 23 (24%) | 57 (32%) |
| > mut |  | 8 (8.5%) | 25 (14%) |
| **gamma** | 272 | 0.48 (-1.22, 3.70) | 0.19 (-2.10, 2.46) |
| **gamma_group** | 272 |  |  |
| > wt |  | 6 (6.4%) | 24 (13%) |
| neutral |  | 47 (50%) | 96 (54%) |
| > mut |  | 41 (44%) | 58 (33%) |
| **MADE** | 272 | 47 (50%) | 82 (46%) |

<sup>†</sup> Statistics presented: Median (Range); n (%)

### *Cis*-regulatory variants contribute significantly to imbalances in the expression of mutant alleles

Next, hypothesizing that both copy number and *cis-*regulatory variants are the major contributors to allelic expression, we set out to assess the contribution of each mechanism towards the net mutant allele expression imbalances detected in these tumors. First, we found positive correlations between overall allelic expression and both copy number and *cis-* regulatory variation (Figure 2C), albeit with an effect for the copy number approximately double the size of that found for *cis-*regulatory variation (average Pearson correlation r-square = 0.82 and 0.39, respectively). Next, considering the variance (Var) of overall allelic expression as the sum of the effects of both mechanisms, plus the covariance (Cov) accounting for predicted non-mutually exclusion of both the mechanisms acting on any given allele:

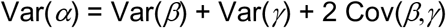

we calculated the contribution of *cis-*regulatory variation to the variance of overall allelic expression as [Var(*γ*) + Cov(*β,γ*)] / Var(*α*). Here, we found that *cis-*regulatory variants explain 17.9% and 16.7% of the variability of overall mutant allelic expression seen in METABRIC and TCGA, respectively.

Finally, assessing how the two mechanisms act simultaneously on each tumor, we found that the majority of samples (71.28% and 54.49% for the METABRIC and TCGA, respectively) had positive γ and negative β values (Figure 2D), meaning that although the mutant allele was in lower genomic quantity, it was nevertheless preferentially expressed compared to the wild-type allele. Only a minor fraction of samples displayed co-occurring preferential allelic expression and a higher allele copy number of the mutant allele (5.32% and 10.11% for the METABRIC and TCGA, respectively). These results were independent of the effect of tumor cellularity (Supplementary Figure 2).

### MADE defines an aggressive subset of *PIK3CA* mutated tumors

To investigate the impact of differential *cis-*regulation of *PIK3CA*’s mutations on clinical outcome (overall and disease-specific survival), we performed univariate survival analysis with γ ratios categorized in two groups: the MADE group (when |γ| ≥ 0.58) and the *mut=wt* group (when |γ| < 0.58). We uncovered that the MADE group had a poorer overall survival rate (*p =* 0.03, Supplementary Figure 3) and disease-specific survival rate (*p =* 0.012, Figure 3A) than the *mut=wt* group for METABRIC. The median overall survival for the MADE group was approximately eight years and for the *mut=wt* group was 18.5 years (Figure 3A), whereas, in the disease-specific analysis, 40% of MADE patients died during the length of the follow-up, in comparison with 20% deaths in the *mut=wt* group (Supplementary Table 4). We further found significantly shorter survival times, in overall and disease-specific survival, for the MADE samples that preferentially expressed the wild-type allele (MADE_wt group, *p =* 0.0073 and =0.0081, respectively) in METABRIC (Figure 3B, Supplementary Figure 3).

**Figure 3:**
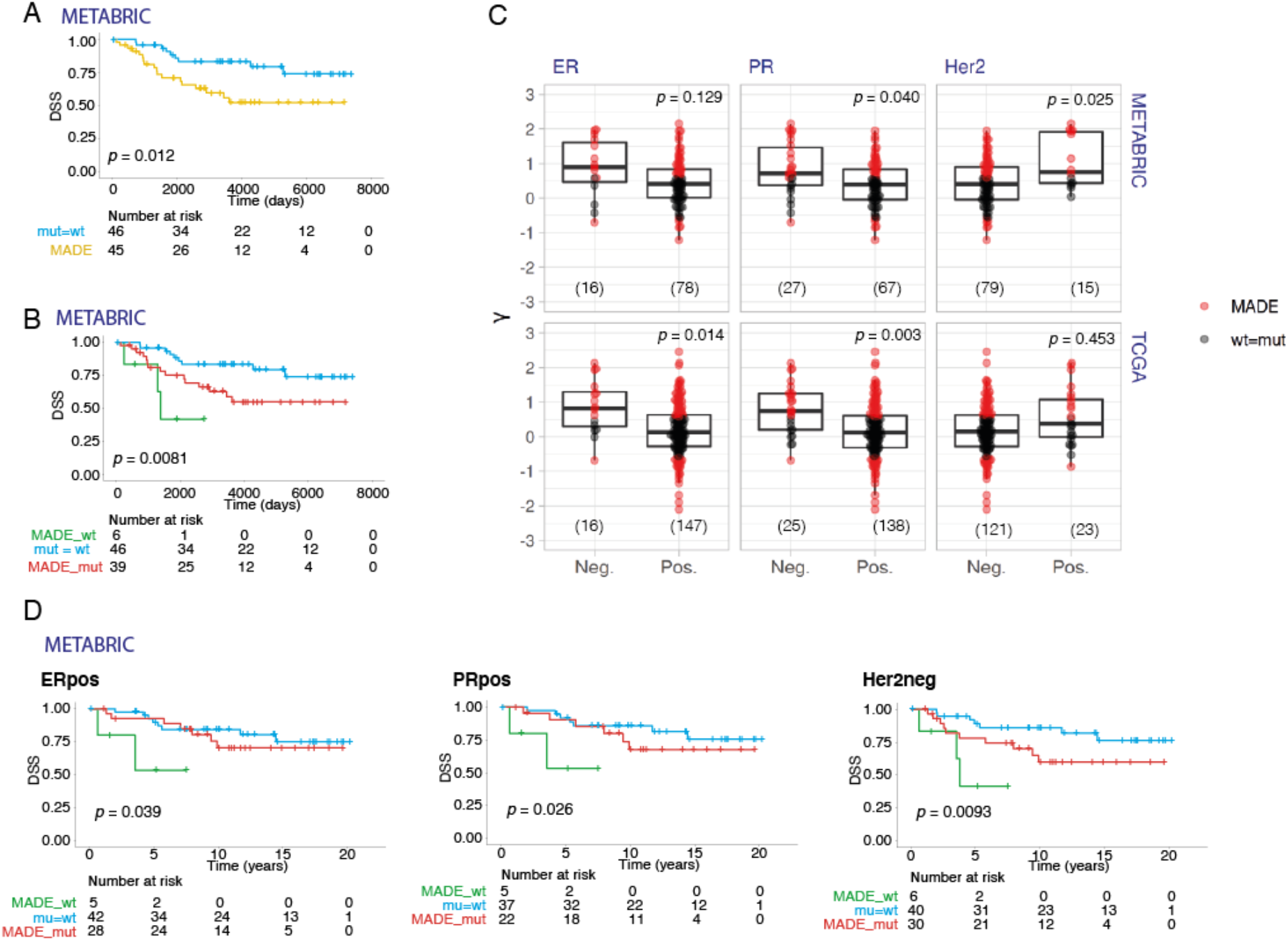
Allelic preferential expression of PIK3CA mutations is associated with survival and clinicopathological parameters in breast cancer. **A –** Kaplan–Meier curves of disease-specific survival showing the worse prognosis of patients with differential expression of the *PIK3CA* mutation (MADE group, shown in yellow) compared to those expressing equimolar levels of mutation and wild-type alleles (*mut=wt* group, shown in blue), in METABRIC. Shown below the graph are the numbers of patients at risk per group throughout time. **B –** Disease-Specific survival Kaplan-Meier curves with the MADE group subdivided into those with preferential expression of the mutant allele (MADE_mut, shown in red) and those with preferential expression of the wild-type allele (MADE_wt, shown in green), confirming worse survival compared to the *mut=wt* group (shown in blue), in METABRIC. Shown below the graph are the numbers of patients at risk per group throughout time. **C** – Preferential expression of the mutated allele is associated with ER-negative, PR-negative, and Her2-positive breast tumors. In all graphs, samples were colored according to the MADE classification. P-values indicated correspond to the Wilcoxon rank sum test with continuity correction, corrected for multiple testing using the Bonferroni correction. **D –** Kaplan-Meier curves showing the worse disease-specific survival of the MADE groups than the *mut=wt* group, in the ER-positive, PR-positive, and Her2-negative subtypes (ER = estrogen; PR = progesterone; Her2 = human epidermal growth factor 2 receptor) in METABRIC.

The categorized γ ratios were not significantly associated with overall survival in the multivariate analysis (Supplementary Figure 4). However, some of the variables that are usually independent prognosis factors, such as PR and Her2 statuses, were not significantly associated with survival either in this analysis. In the TCGA set, there was no significant difference in overall survival for either α or γ (Supplementary Figure 5). However, due to the relatively shorter follow-up time of this data set (median ~1 year) and the fact that tumors were mainly Luminal A (~61.2% of samples) [15], the power to detect significant differences is smaller than that of METABRIC.

### *PIK3CA* MADE associates with clinicopathological variables

Next, we sought to investigate whether *PIK3CA*’s MADE was associated with known prognostic clinicopathological variables, namely hormone receptors (ER, PR) and Her2 amplification (Figure 3C, Table 3), which are directly and indirectly connected to gene expression regulation, respectively.

**Table 3.** Statistical analysis between normalized and raw mutant allele expression ratios. P-values indicated correspond to two-sided Wilcoxon rank sum tests with continuity correction, adjusted per study with the Bonferroni method.

|  | METABRIC |  |  | TCGA |  |  |
| --- | --- | --- | --- | --- | --- | --- |
| | n (pos/neg) | $\alpha$ | $\gamma$ | n (pos/neg) | $\alpha$ | $\gamma$ |
| *Clinical Features* |  |  |  |  |  |  |
| ERα status | 78/16 | 0.026 | 0.129 | 147/16 | 0.157 | 0.014 |
| PR status | 67/27 | 0.007 | 0.040 | 138/25 | 0.054 | 0.003 |
| Her2 status | 15/79 | 0.018 | 0.025 | 23/121 | 1.000 | 0.453 |

Legend: ER = estrogen; PR = progesterone; Her2 = human epidermal growth factor 2 receptor

For METABRIC, the mean α was significantly higher in ER-negative tumors (*p =* 0.026), PR-negative tumors (*p =* 0.007) and Her2-positive tumors (*p =* 0.018) (Supplementary Figure 6). When evaluating the contribution of *cis-*regulatory variation to this association, we also found that higher average γ values associated with lower PR expression (*p =* 0.040) and Her2-positive tumors (*p =* 0.025), but we did not find a significant association with ER expression (*p =* 0.129) (Figure 3C). These results showed that the predominant abundance of the mutant allele was associated with worse prognosis variables (ER negativity, PR negativity, Her2positivity), particularly in tumors with preferential mutant allele expression. The analysis of the TCGA set corroborated the significant association between the PR status and γ ratios (*p =* 0.003) (Figure 3C, Supplementary Figure 6). However, it showed an association between ER status and γ values only (*p =* 0.014), albeit not with Her2 status.

Given these results, we took MADE into consideration in the survival analysis within the expression subgroups of ER, PR, and Her2. Both MADE vs. *mut=wt* groups did not reveal significant differences in overall and disease-specific survival. However, there was a trend for worse survival for the MADE category in the Her2 negative subgroup (*p =* 0.056 and 0.027 in overall and disease-specific analyses, respectively) (Supplementary Figure 3C). However, when further stratifying the MADE groups, a poorer prognosis for MADE_wt samples was identified in ER-positive (*p =* 0.004), PR positive (*p =* 0.0013) and Her2 negative tumors (*p =* 0. 0066) (Figure 3D). Thus, MADE defines an aggressive subgroup within the usually good prognosis ER-positive, PR-positive, and Her2-negative subsets of tumors.

Considering other known prognostic variables, including tumor size, grade, and molecular subtypes (PAM50 [16] and IntClust [14]), we found a significant association between γ ratios and PAM50 subtypes only in TCGA (*p =* 0.002) (Supplementary Figure 7, Supplementary Table 5).

Finally, we did not find an association between the candidate germline regulatory variant rs2699887 and MADE or clinical outcome, suggesting that somatic variants are most likely involved in the significant associations described above (Supplementary Figure 8).

## Discussion

Our work reveals the role of *cis-*regulatory variation acting on *PIK3CA* somatic mutations as modifiers of mutation penetrance. We show for the first time that allelic expression imbalance between *PIK3CA’s* mutant and wild-type alleles is common and prognostic in breast cancer. Particularly, preferential expression of the mutant allele is significantly more common than expected, and whether there is a positive selection for it in tumor evolution should be further investigated. However, the genomic allelic imbalance remains the determinant of allelic expression dosage with a larger effect, with *cis-*regulatory variation responsible for ~18% of the variability in overall allelic expression in tumors. The analysis of RNA-seq data from two independent cohorts of tumor samples, the METABRIC and TCGA projects, strongly supports our findings.

Moreover, we show that preferential expression of the mutant allele is associated with poor prognosis variables, such as PR [19] negative and Her2-positive [20] statuses. In the METABRIC dataset, we also found that patients with tumors with mutant allelic expression imbalance had worse overall survival than those without it, particularly in the ER-positive, PR-positive, and Her2-negative subsets. Interestingly, the group with preferential expression of the wild-type allele showed the worst outcome. In these tumors, *PIK3CA* mutations are lowly expressed and are possibly passenger events. Hence, the poor prognosis might be due to other mutations or combinations thereof. Albeit the significant findings, one caveat of our study is the minimal size of this subgroup of samples. Identifying what other events might be contributing to the worse survival requires a larger sample set and combinatorial analysis with additional data on other mutations.

Besides the potential use of our findings as a prognosis biomarker in the clinic, these results may also have therapeutic implications. Some of the major clinical challenges in cancer treatment are identifying biomarkers of prognosis and defining which patients will benefit from a given therapy. In particular, identifying patients unlikely to respond to specific therapies is crucial to prevent unnecessary drug cytotoxicity without any therapeutic benefits. Our results reveal the importance of considering allelic expression in somatic mutation screens in these two aspects of patient management. Despite the high frequency of *PIK3CA* mutation in breast cancers, the response to PI3K inhibitor therapy has been unsuccessful as expected, and the prognostic significance of detecting somatic *PIK3CA* mutations in breast tumors is unclear [21]. Relevant to this discussion, we have previously shown that the presence of *PIK3CA* mutations confer a poorer prognosis in patients with ER-positive breast cancer only when stratified into copy-number driven subgroups (IntClust 1+,2+,9+) [22]. In the present study, we provide new evidence for the prognostic significance of these mutations at the expression level in unstratified ER-positive tumors. This prognostic significance has a potential impact on therapy response and clinical management since one may hypothesize that little to no benefit would come from treatment in the cases not expressing the targetable mutation. Further studies evaluating the allelic expression of mutant oncogenes in the tumors of patients enrolled in molecular-driven trials will clarify this impact.

More challenging is determining which *cis-*regulatory mechanisms are promoting allelic expression imbalances. Both inherited [10,23] and acquired variants [6,24–26], and epigenetics can affect gene expression in an allelic manner [27,28]. Here we show that normal *cis-*regulatory variation controls *PIK3CA’s* expression in normal breast tissue, and our data suggest that the SNP rs2699887 is possibly the main regulatory variant. We believe the mechanism includes modification of an NF-YA transcription factor binding site at the promoter of *PIK3CA*, supported by *in-silico* and *in-vitro* data. We also found that the heterozygotes were associated with higher expression of the *PIK3CA* gene than the common homozygotes. Although there is published data supporting the clinical association of rs2699887 with poor prognosis in other cancers [29,30], linked to an increase in PI3K signaling, there is still some data supporting the opposite association [31,32]. We did not find an association between rs2699887 and survival, suggesting that other mechanisms besides normal *cis-*regulatory variation contribute to the preferential allelic expression in these tumors. Double *PIK3CA* mutations in the same allele are frequent in breast tumors [33], and the impact of non-coding mutations in cancer is just starting to be explored [6]. So, a strong possibility is that the combination of non-coding and coding mutations in the same gene might be underlying the allelic expression imbalances we are detecting.

Further studies on allelic expression imbalances of activating mutations, and even inactivating ones, should further reveal the contribution of *cis-*regulatory mechanisms in tumor development and progression. Particularly interesting to determine is whether the coding mutation originates in an allele predisposed with higher expression, or whether a sequence of somatic events introduces the coding activating mutation and additional *cis-*regulatory non-coding mutations. The answers could have significant repercussions on our understanding of tumor evolution.

## Conclusion

In summary, we show that differential expression between the mutant and wild-type alleles of *PIK3CA* is common in breast cancer and mainly due to allele-specific *cis*-regulatory effects. We further show that mutant allele differential expression is associated with clinical parameters such as ER, PR, and Her-2 statuses and is prognostically significant. Collectively, our work establishes the prognostic relevance of allele-specific transcriptional regulation of somatic mutations. It also supports a shift in the mutation testing in patient management, where the level of expression of mutations should be considered, besides the detection at the DNA level.

## Methods

### Differential allelic expression analysis

DNA and total RNA from 64 samples of normal breast tissue were hybridized onto Illumina Exon510S-Duo arrays (humanexon510s-duo), and data were analyzed as described before [17]. In short, after sample filtering and normalization, SNPs with average RNA log_2_ allelic intensity values greater than 9.5 and heterozygous in five or more samples were kept for further analysis. Allelic log-ratios were calculated for RNA and DNA intensity data: log-ratio = log_2_ A −log_2_ B, for alleles A and B. A two-sample Student’s t-test was applied to compare RNA log-ratios between heterozygous (AB) and homozygous groups (AA and BB). Only SNPs with p-values lower than 0.05 for all comparisons were further analyzed. Allelic expression (AE) ratios were normalized for allelic DNA content (RNA log-ratio – DNA log-ratio), and differential allelic expression (DAE) defined as |AE ratio| ≥ 0.58 (1.5-fold or greater between alleles). This threshold was established based on the sensitivity and specificity of the applied DAE detection method. Linkage disequilibrium (LD) between daeSNPs was evaluated using the genetic variant-centered annotation browser SNiPA [34].

### Genotype imputation analysis on normal breast tissue samples

Genotype imputation was run on the Illumina Exon 510 Duo germline genotype data from the 64 samples that passed microarrays quality control filters. Before imputation, genotyping data were quality controlled, and SNPs with call rates < 85%, minor allele frequency < 0.01, and Hardy-Weinberg equilibrium with *p =* < 1.0e^−05^ were excluded from the analysis. Genotypes were imputed with MACH1.0 [35] for all additional known SNPs on chromosome 3 from good quality Illumina SNP genotype. The phased haplotypes from the HapMap3 release (HapMap3 NCBI Build 36, CEU panel - Utah residents with Northern and Western European ancestry) were used as the reference panel. For imputation with MaCH1.0, we applied the recommended two-step imputation process: model parameters (crossover and error rates) were estimated before imputation using all haplotypes from the study subjects and running 100 Hidden Markov Model (HMM) iterations with the command options: - greedy and -r 100. Genotypes were imputed using the model parameter estimates from the previous round with options: -greedy, -mle, and -mldetails. Imputation results were filtered based on an rq score higher than 0.3 [35], a platform-specific measurement of SNP imputation uncertainty.

### Differential allelic expression (DAE) mapping analysis on normal breast tissue samples

Differential allelic expression mapping analysis was performed by stratifying AE ratios at each *PIK3CA* daeSNP according to the genotype at the genotyped/imputed SNPs located within ±250Kb of each daeSNP. A two-sample t-test was applied to assess differences between the mean AE ratio between the heterozygous group samples and the combined homozygous groups. Permutation-corrected p-values (1000 permutations) were considered significant below 0.05 for variants with heterozygous samples that displayed greater mean fold differences between alleles than homozygous samples.

Proxy SNPs were identified using HaploReg v4.1 available at https://pubs.broadinstitute.org/mammals/haploreg/haploreg.php, using genotype data from the Phase 1 of the 1000G Project, for the CEU population, with an LD threshold of r^2^ ≥0.8 [36,37].

### Functional annotation of associated variants

Variants associated with AE levels were assessed for overlap with epigenetic marks derived from the Encyclopedia of DNA Elements (ENCODE) and NIH Roadmap Epigenomics projects, such as chromatin states (chromHMM) annotation, regions of *DNase* I hypersensitivity, transcription factor binding sites, and histone modifications of epigenetic markers (H3K4Me1, H3K4Me3, and H3K27Ac) (http://genome.ucsc.edu/ENCODE/) for normal human mammary epithelial cells (HMECs), human mammary fibroblasts (HMFs), BR.MYO (breast myoepithelial cells) and BR.H35 (breast vHMEC) and two breast cancer cell lines MCF-7 and T47D. We prioritized variants located on either active promoter or enhancer regions in mammary cell lines, and for which ChIP-Seq data indicated protein binding or position weight matrix (PWM) scores predicted differential protein binding for different alleles. Two publicly available tools, RegulomeDB and HaploReg v4.1 and the MotifBreakR Bioconductor package, were also used to evaluate those candidate functional variants [36–38].

### Electrophoretic mobility shift assay (EMSA)

MCF-7 (ER-positive) and HCC1954 (ER-negative) breast cancer cell lines were cultured in DMEM and RPMI culture media, respectively, supplemented with 10% FBS and 1% PS (penicillin and streptomycin). Nuclear protein extracts were prepared using the Thermo Scientific Pierce^™^ NER kit, according to the manufacturer’s instructions. Oligonucleotide sequences corresponding to the C (common) and T (minor) alleles of rs2699887 (5’-AGCGTGAGTAGAGCGCGGA[C/T]TGGCCGGTAGCGGGTGCGGTG-3’) were labeled using the Thermo Scientific Pierce Biotin 3’ End DNA Labelling Kit, according to the manufacturer’s instructions. Antibodies used for supershift competition assays were NF-YA (H-209) (Santa Cruz Biotechnology, SC-10779X) and HMGA1a/HMGA1b (Abcam, ab4078). EMSA experiments were performed using the Thermo Scientific LightShift^™^ Chemiluminescent EMSA Kit, using the buffer and binding reaction conditions previously described [39]. Each EMSA was repeated at least twice for all combinations of cell extract and oligonucleotide, which were also tested in serial dilution amounts.

### Breast tumor samples

The METABRIC dataset of tumor samples included 2433 samples from the METABRIC project [14] with DNA sequencing data, among which 480 were subjected to a capture-based RNA sequencing study. Sequencing libraries were generated as previously described. In brief, sequencing libraries using total RNA generated from frozen tissues with a TruSeq mRNA Library Preparation Kit using poly-A-enriched RNA (Illumina, San Diego, CA, USA) and enriched with the human kinome DNA capture baits (Agilent Technologies, Santa Clara, CA, USA). Six libraries were pooled for each capture reaction, with 100 ng of each library and sequenced (paired-end 51bp) on an Illumina HiSeq2000 platform. We selected a subset of samples with DNA and RNA sequencing data and *PIK3CA* missense mutations for further analysis.

The TCGA dataset comprised 695 samples from TCGA breast cancers [40], from which we selected a subset of 289 samples with *PIK3CA* missense mutations for further analysis. Supplementary Table 2 summarizes the demographic features and disease characteristics of the two datasets.

### DNA-seq and RNA-seq variant calling in tumors

#### Alignment & Preprocessing

Sequence data (FASTQ) mapped to reference genome (hg19) were aligned using STAR v2.4.1 [41]. A two-pass alignment was carried out: splice junctions detected in the first alignment run are used to guide the final alignment. Duplicates were marked with Picard v1.131 (http://picard.sourceforge.net). Genome Analysis Toolkit (GATK) was used for indel realignment and base quality score recalibration [42].

#### Variant Calling and Annotations

SNV and indel variants were called using GATK Haplotype Caller. Hard filters using GATK VariantFiltration were applied to variants [42]. Variants were annotated with Ensembl Variant Effect Predictor (VEP) [43]

Heterozygous genotypes were called from DNA data to avoid RNA editing and other RNA related variants because true allelic imbalance can lead to heterozygous sites being called homozygous in RNA-based genotype calling.

#### Statistical analysis of allelic expression imbalances in tumors

Before the analysis, a set of filtering steps was performed to select samples: 1) good quality data; 2) presence of missense mutations; 3) and a minimum of 30 reads for RNA and DNA data. Clinical data for METABRIC were updated from the original studies with the latest available records. Clinical data for TCGA were imported from https://portal.gdc.cancer.gov/ on 26/11/2018.

For all samples, three parameters were calculated: 1) The DNA mutant allele ratio β = Log_2_[(mutant allele read count in DNA)/(wild-type allele read count in DNA)], which served to control for sequencing artifacts from heterozygous genotypes and to account for differences in variant frequencies in DNA; 2) α = Log_2_[(mutant allele read count in RNA)/(wild-type allele read count in RNA)], that served as a measure of the net allelic expression imbalance in tumors; and 3) the normalized mutant allele expression ratio γ = α − β, which is a measure of mutant allelic expression imbalance due to *cis-*regulation alone.

Association between allelic expression imbalance ratios and clinical data was achieved by bivariate analysis Wilcoxon rank sum test with continuity correction or Kruskal-Wallis rank sum test, as indicated in tables and figures. P-values were adjusted per study using the Bonferroni correction and were considered significant when ≤ 0.05. Correlation analysis “alpha vs. beta” and “alpha vs. gamma” ratios for both sets of samples were performed using a Pearson’s test. All statistical analysis and data visualization were performed using R.

Additionally, we defined mutant allele differential expression (MADE) as |γ|≥ 0.58 (1.5-fold or larger difference). In the survival analysis, we further separated the MADE samples into MADE_wt (γ ≤ −0.58, or preferential expression of wild-type allele) and MADE_mut (γ ≥ 0.58, or preferential expression of mutant allele).

#### Survival Analyses

Kaplan-Meier plots and multivariate Cox proportional hazard models were used to examine the association between alpha and gamma allelic expression ratios and survival using the *survival* package from R [44,45]. Death due to all causes was used as the endpoint, and all alive subjects were censored at the date of the last contact. Kaplan-Meier survival curves were compared using the log-rank test.

For the multivariate analysis, Cox proportional hazard model was used to assess the effect of γ on the overall survival. Hazard ratios (HRs) and 95% confidence intervals (CI) were estimated by fitting the Cox model while adjusting for age and tumor characteristics, such as size, Scarff-Bloom-Richardson histological grade, clinical stage and estrogen receptor (ER), progesterone (PR), and human epidermal growth factor 2 (Her2) statuses. All t-tests were two-sided, and *p* ≤ 0.05 were considered statistically significant.

## Declarations

### Ethics approval and consent to participate

Normal breast and tumor samples were obtained with the consent from donors and appropriate approval from local ethical committees, with the detailed information described in the respective original publications: normal tissue [8], METABRIC [14], TCGA [40].

### Availability of Data and Code

Microarray raw data are deposited in the Gene Expression Omnibus under accession number GSE35023. Primary data (BAM files) for DNAseq are deposited at the European Genome-phenome Archive (EGA) under study accession number EGAS00001001753 and may be downloaded upon request and authorization by the METABRIC Data Access Committee. Primary data (BAM files) for RNAseq are currently being deposited in EGA (accession number to be determined). Primary data (BAM files) for DNAseq and RNAseq from TCGA are deposited in the database of Genotypes and Phenotypes (dbGaP) under the study accession number phs000178. The filtered data and code for the analysis of mutant allele expression imbalances and the survival analysis can be publicly accessed at https://github.com/maialab/pik3ca.

### Competing interests

The authors declare that they have no competing interests.

### Funding

This work was supported by Portuguese national funding through FCT-Fundação para a Ciência e a Tecnologia, and CRESC ALGARVE 2020, institutional support ALG-01-0145-FEDER-31477 – “DevoCancer”, CBMR - UID/BIM/04773/2013, POCI-01-0145-FEDER-022184 - ”GenomePT”, the contract DL 57/2016/CP1361/CT0042 (J.M.X.) and individual fellowships SFRH/BPD/99502/2014 (J.M.X.) and PD/BD/114252/2016 (F.E.). Funding was also received from the People Programme (Marie Curie Actions) of the European Union’s Seventh Framework Programme FP7/2007-2013/ under REA grant agreement n° 303745 (ATM), and a Maratona da Saúde Award (ATM).

The METABRIC project was funded by Cancer Research UK, the British Columbia Cancer Foundation, and the Canadian Breast Cancer Foundation BC/Yukon. This sequencing project was funded by CRUK grant C507/A16278 and CRUK core grant A16942. The authors also acknowledge the support of the University of Cambridge, Hutchinson Whampoa, the NIHR Cambridge Biomedical Research Centre, the Cambridge Experimental Cancer Medicine Centre, the Centre for Translational Genomics (CTAG) Vancouver, and the BCCA Breast Cancer Outcomes Unit. We thank the Genomics, Histopathology, and Biorepository Core Facilities at the Cancer Research UK Cambridge Institute and the Addenbrooke’s Human Research Tissue Bank (supported by the National Institute for Health Research Cambridge Biomedical Research Centre).

### Authors’ Contributions

L.C., J.M.X., S.F.C., and A.T.M. wrote the manuscript.

J.M.X., S.F.C., and A.T.M. contributed to the overall design of this study.

Data were collected by F.E., A.B., L.M., R.B., C.C., S.F.C., and A.T.M.

Data were analyzed and interpreted by L.C., J.M.X., R.M., B.P.A., F.E., C.S., I.D., M.E, A.M., I.A.S., J.S., and A.T.M.

All authors have read and approved the final version of the manuscript.

## Acknowledgments

We thank all the patients who donated tissue and the associated pseudo-anonymized clinical data for this project. The authors would also like to thank the Functional Genomics of Cancer group members at CINTESIS-UAlg for helpful discussions and Vitor Morais at UAIC for administrative support.

## Notes

https://github.com/maialab/pik3ca

### Competing Interest Statement

The authors have declared no competing interest.

### Summary of Updates

Updated title, abstract, and introduction for clarity. Addition of a supplementary figure. Authors' affiliations updated.

