## Supplementary Figures for "Allelic expression imbalance of *PIK3CA* mutations is frequent in breast cancer and prognostically significant"

Lizelle Correia et al.

### Supplementary Figures

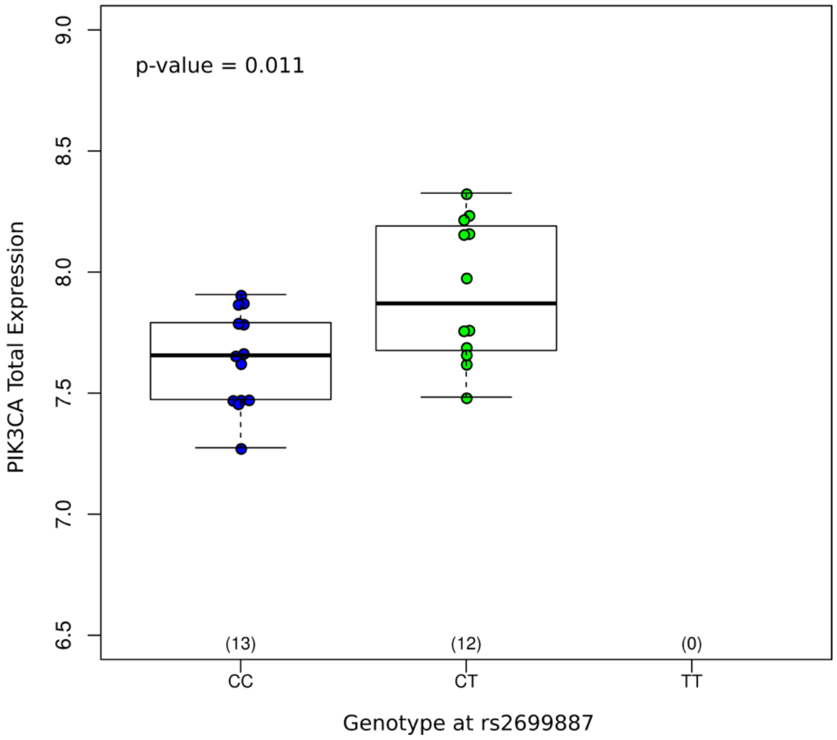


**Supplementary Figure 1 – rs2699887 is an eQTL for the expression of *PIK3CA* in tumors from METABRIC.** P-value indicated corresponds to Student's t-test.

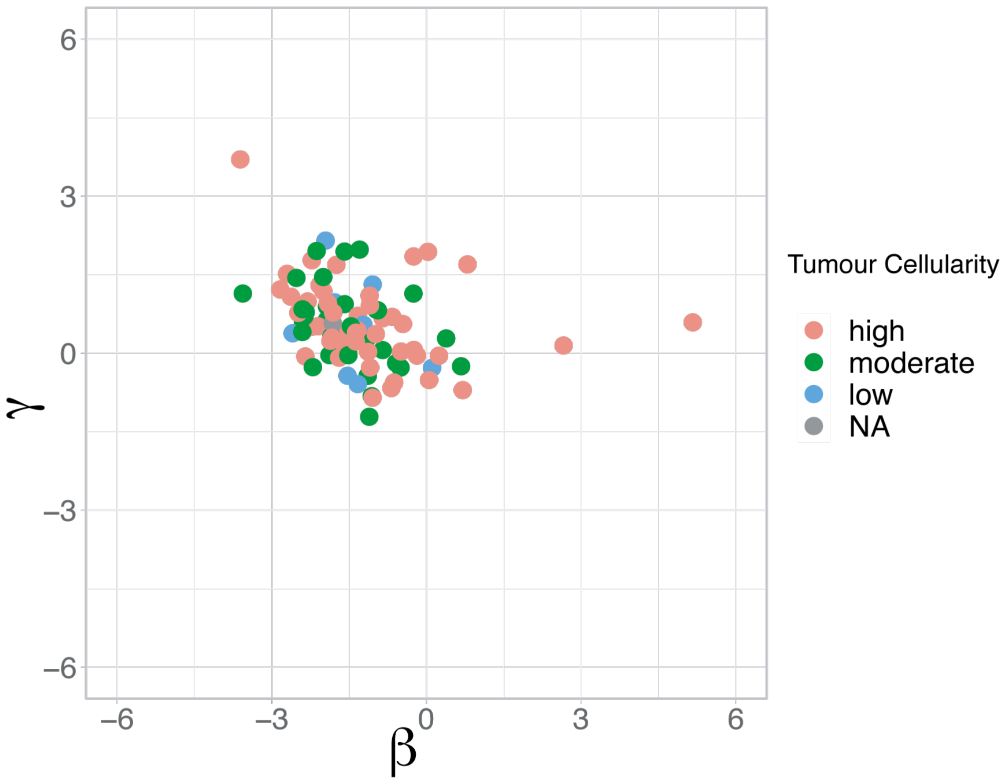


**Supplementary Figure 2 - Comparison of *PIK3CA's* matched β and γ values reveals no pattern of association between ratios and cellularity in METABRIC tumors.** Each dot represents a tumor, and color indicates cellularity levels.

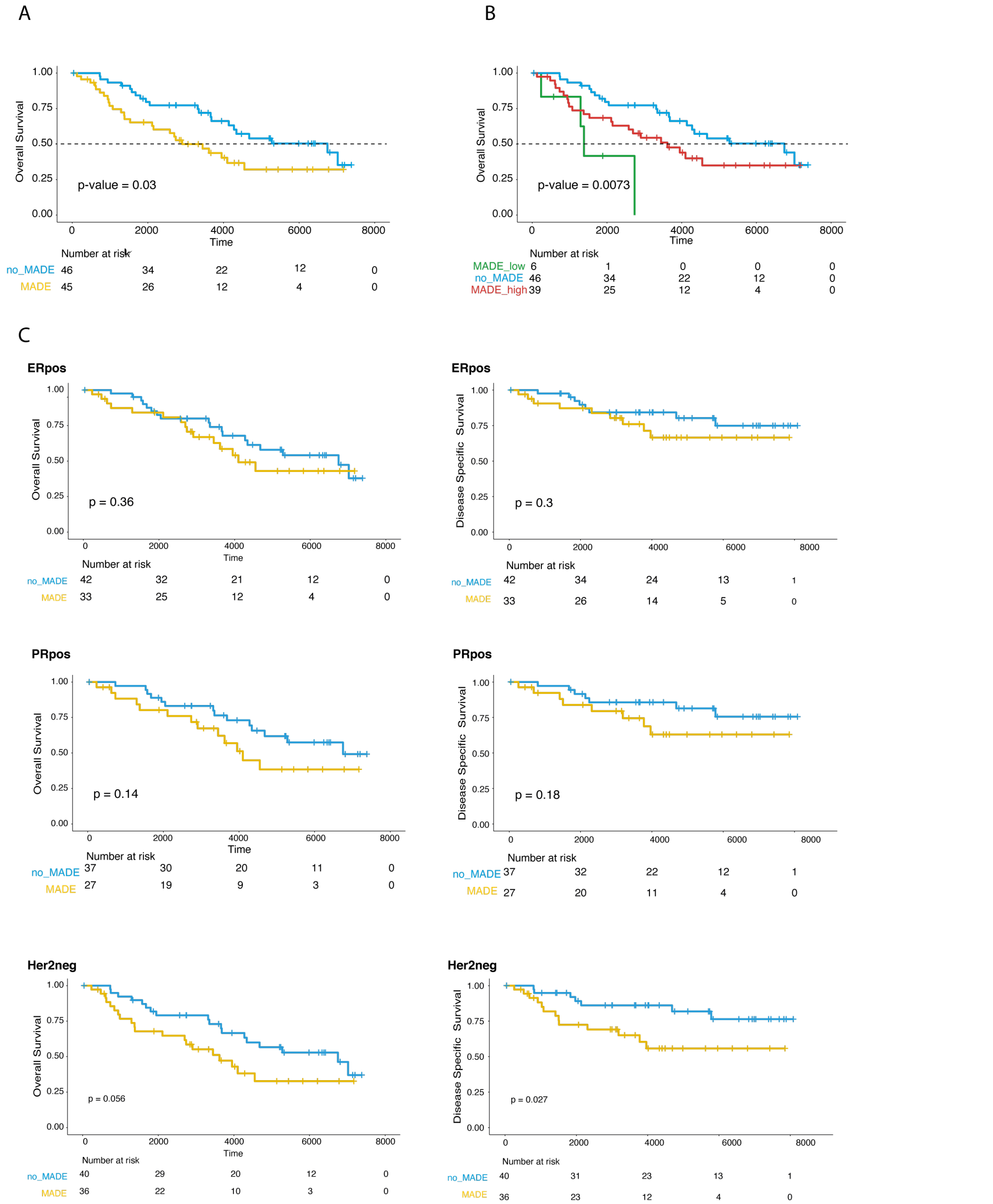


**Supplementary Figure 3 - Survival of mutant allele preferential expression in METABRIC tumors. A** – Kaplan–Meier curves of overall survival showing the worse prognosis of patients with differential expression of the *PIK3CA* mutation (MADE group, shown in yellow) compared to those expressing equimolar levels of mutation and wild-type alleles (mut=wt group, shown in blue). Shown below the graph are the numbers of patients at risk per group throughout time. **B** – Overall survival Kaplan-Meier curves with the MADE group subdivided into those with preferential expression of the mutant allele (MADE_mut, shown in red) and those with preferential expression of the wild-type allele (MADE_wt, shown in green), confirming worse survival compared to the mut=wt group. Shown below the graph are the numbers of patients at risk per group throughout time. **C** –Kaplan-Meier curves showing the worse overall survival of the MADE groups than the mut=wt group, by in the ER-positive, PR positive, and Her2 negative subtypes (ER = estrogen; PR = progesterone; Her2 = human epidermal growth factor 2 receptor).

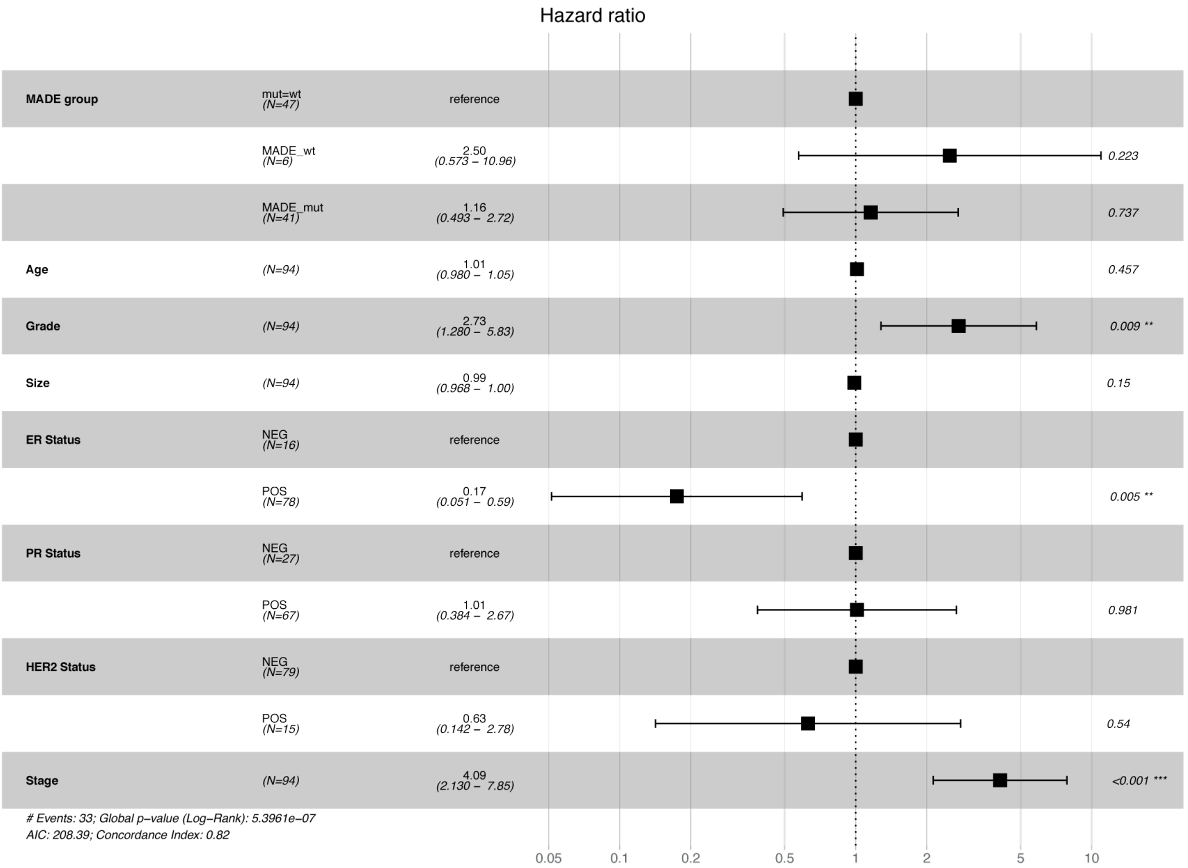


**Supplementary Figure 4 - Survival of mutant allele preferential expression in METABRIC tumors.** Forest plot from multivariate Cox regression model showing the association of PIK3CA's γ ratios and overall survival, adjusted for age, grade, ER, PR and HER2 statuses, and tumor stage. (ER = estrogen; PR = progesterone; Her2 = human epidermal growth factor 2 receptor).

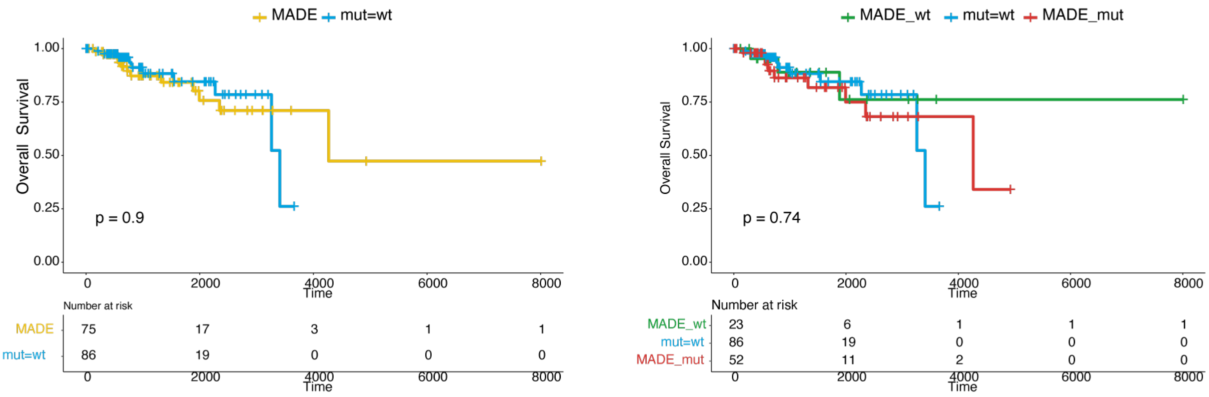


**Supplementary Figure 5 - Survival analysis of mutant allele preferential expression in TCGA tumors. A** – Kaplan–Meier curves of overall survival showing no difference in survival between patients with differential expression of the *PIK3CA* mutation (MADE group, shown in yellow) and those expressing equimolar levels of mutation and wild-type alleles (mut=wt group, shown in blue). Shown below the graph are the numbers of patients at risk per group throughout time. **B** – Overall survival Kaplan-Meier curves with the MADE group subdivided into those with preferential expression of the mutant allele (MADE_mut, shown in red) and those with preferential expression of the wild-type allele (MADE_wt, shown in green), confirming no difference in survival compared to the mut=wt group. Shown below the graph are the numbers of patients at risk per group throughout time.

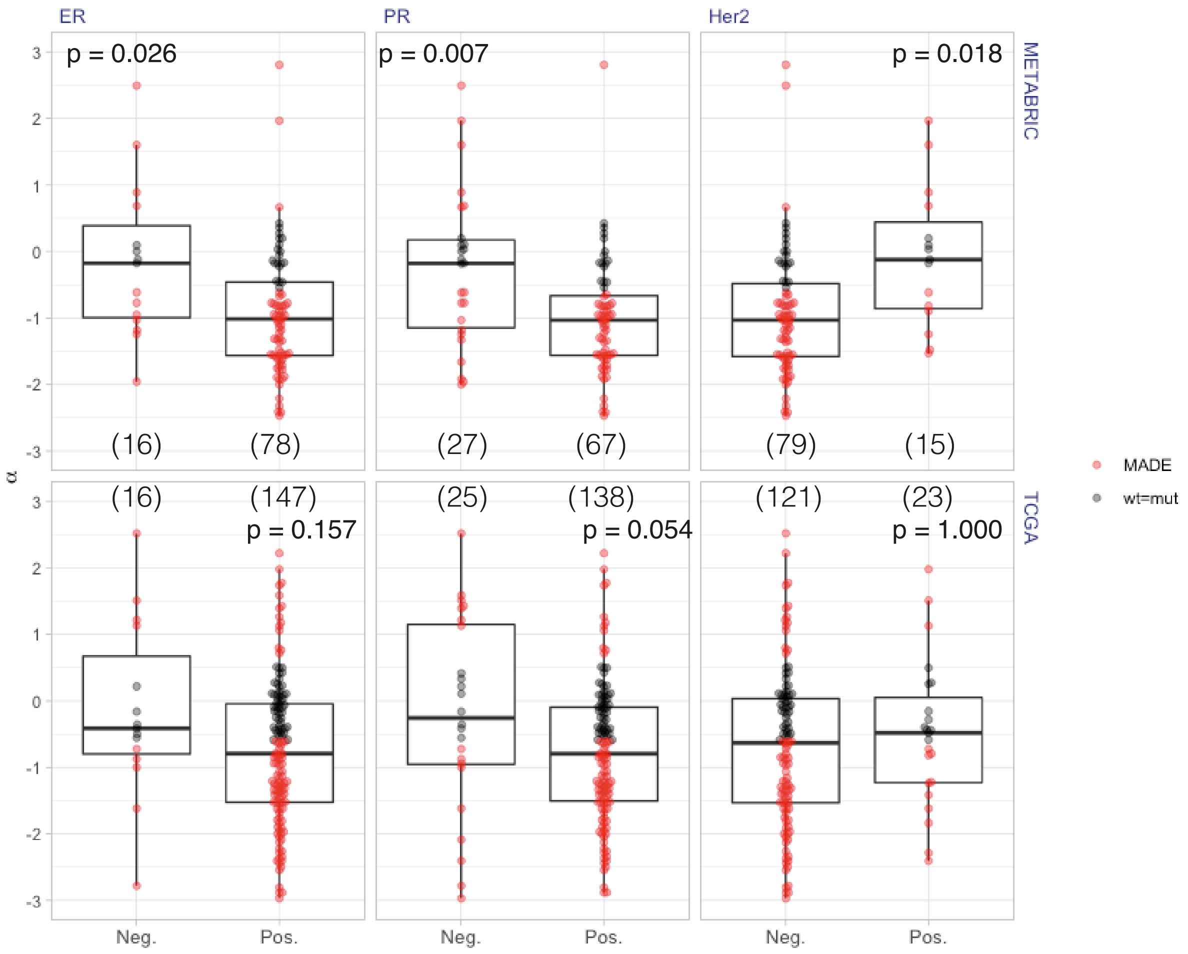


**Supplementary Figure 6 - Association analysis between α ratios and ER, PR, and HER2 statuses.** Boxplots for α ratios for the METABRIC (top) and TCGA (bottom) tumors. Each dot represents a sample, and the numbers of samples in each group are indicated in brackets. In all graphs, samples were colored according to the MADE classification. P-values indicated correspond to the Wilcoxon rank sum test with continuity correction, corrected for multiple testing using the Bonferroni correction. (ER = estrogen; PR = progesterone; Her2 = human epidermal growth factor 2 receptor).

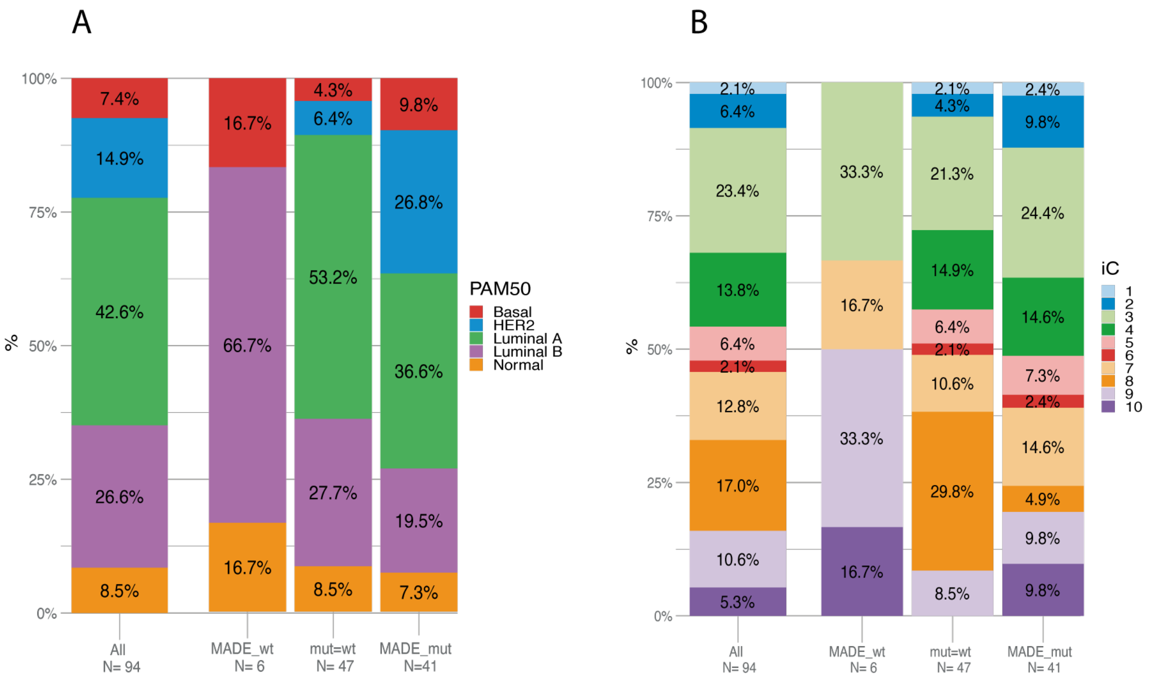


**Supplementary Figure 7 - Distribution (A) PAM50 (Prediction Analysis of Microarray 50) subtypes and (B) IntClust (Integrative Cluster Classification) within *PIK3CA's* MADE groups in METABRIC set.**

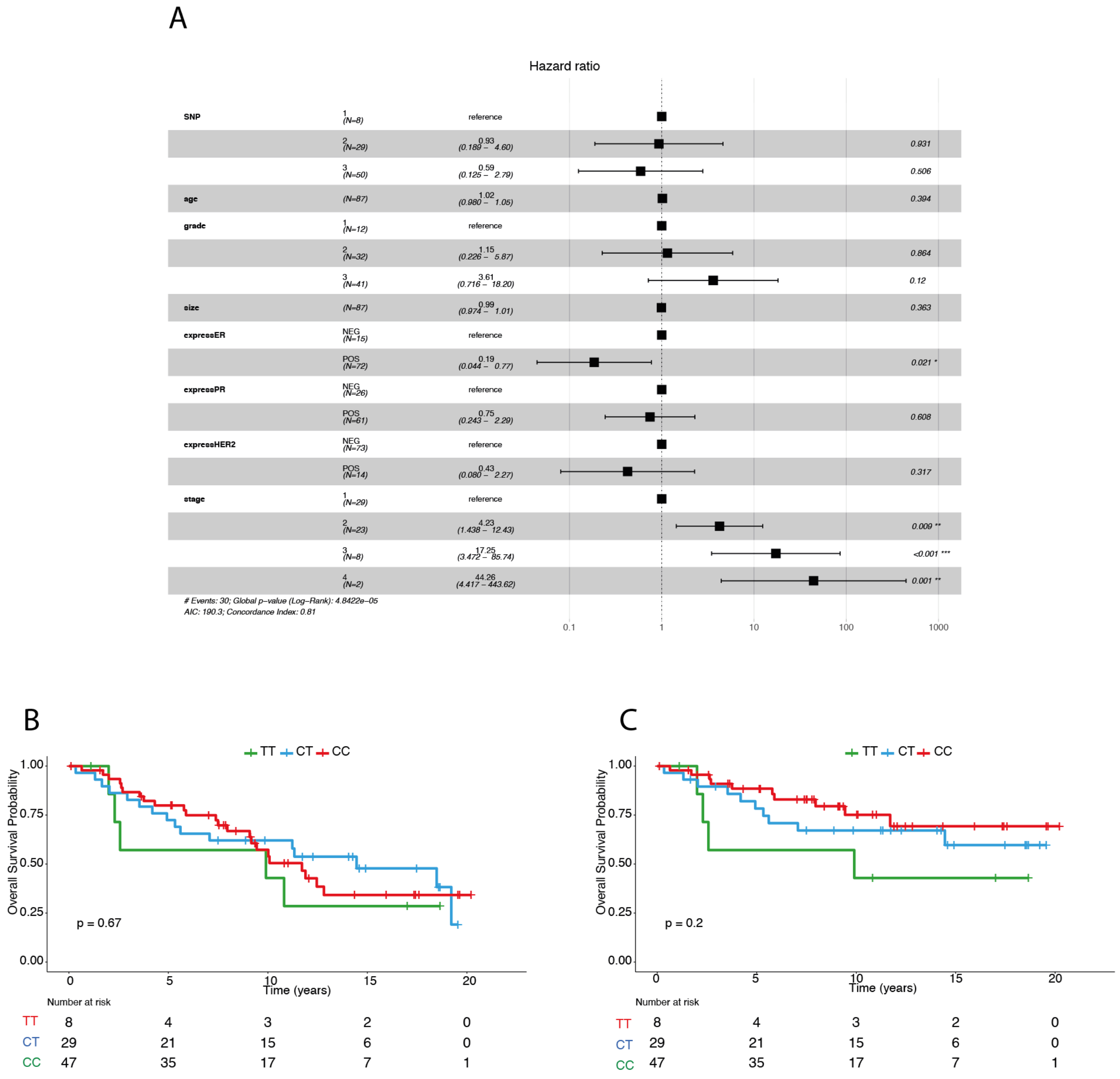


**Supplementary Figure 8 - Survival analysis of genotype at rs2699887 in METABRIC tumors. A -** Forest plot from multivariate Cox regression model showing the association of genotype at rs2699887 and overall survival, adjusted for age, grade, ER, PR and HER2 statuses, and tumor stage. (ER = estrogen; PR = progesterone; Her2 = human epidermal growth factor 2 receptor). **B** and **C -** Kaplan–Meier curves of overall and disease-specific survival, respectively, showing no association between genotype at rs2699887 and survival in METABRIC. Shown below the graph are the numbers of patients at risk per group throughout time.
