## Supplementary Tables for "Allelic expression imbalance of *PIK3CA* mutations is frequent in breast cancer and prognostically significant"

Lizelle Correia et al.

**Supplementary Table 1:** Linkage disequilibrium between daeSNPs of *PIK3CA*. Data was mapped and annotated according to the “grch37/1kgpp3v5” release, European population, using SNiPA (https://snipa.helmholtz-muenchen.de).

| QRSID | RSID | R2 | DPRIME |
|---|---|---|---|
| **rs7636454** | rs4855093 | 0.91 | 0.97 |
| **rs7636454** | rs3960984 | 0.90 | 0.98 |
| **rs7636454** | rs9838411 | 0.91 | 0.98 |
| **rs7636454** | rs3729679 | 0.22 | 1.00 |
| **rs7636454** | rs12488074 | 0.99 | 1.00 |
| **rs7636454** | rs7636454 | 1.00 | 1.00 |
| **rs3960984** | rs4855093 | 0.98 | 1.00 |
| **rs3960984** | rs3960984 | 1.00 | 1.00 |
| **rs3960984** | rs9838411 | 0.99 | 1.00 |
| **rs3960984** | rs3729679 | 0.20 | 0.99 |
| **rs3960984** | rs12488074 | 0.89 | 0.98 |
| **rs3960984** | rs7636454 | 0.90 | 0.98 |
| **rs3729679** | rs4855093 | 0.20 | 0.98 |
| **rs3729679** | rs3960984 | 0.20 | 0.99 |
| **rs3729679** | rs9838411 | 0.21 | 0.99 |
| **rs3729679** | rs3729679 | 1.00 | 1.00 |
| **rs3729679** | rs12488074 | 0.22 | 1.00 |
| **rs3729679** | rs7636454 | 0.22 | 1.00 |
| **rs12488074** | rs4855093 | 0.90 | 0.97 |
| **rs12488074** | rs3960984 | 0.89 | 0.98 |
| **rs12488074** | rs9838411 | 0.91 | 0.98 |
| **rs12488074** | rs3729679 | 0.22 | 1.00 |
| **rs12488074** | rs12488074 | 1.00 | 1.00 |
| **rs12488074** | rs7636454 | 0.99 | 1.00 |
| **rs4855093** | rs4855093 | 1.00 | 1.00 |
| **rs4855093** | rs3960984 | 0.98 | 1.00 |
| **rs4855093** | rs9838411 | 0.99 | 1.00 |
| **rs4855093** | rs3729679 | 0.20 | 0.98 |
| **rs4855093** | rs12488074 | 0.90 | 0.97 |
| **rs4855093** | rs7636454 | 0.91 | 0.97 |
| **rs9838411** | rs4855093 | 0.99 | 1.00 |
| **rs9838411** | rs3960984 | 0.99 | 1.00 |
| **rs9838411** | rs9838411 | 1.00 | 1.00 |
| **rs9838411** | rs3729679 | 0.21 | 0.99 |
| **rs9838411** | rs12488074 | 0.91 | 0.98 |
| **rs9838411** | rs7636454 | 0.91 | 0.98 |

**Supplementary Table 2:** Candidates SNPs from mapping analysis of DAE data in *PIK3CA*.

| daeSNP | SNP associated | Perm pvalue |
|---|---|---|
| rs3960984 | rs4854984 | 0.001 |
| rs3960984 | rs1468920 | 0.001 |
| rs3960984 | rs1468922 | 0.009 |
| rs3960984 | rs1468923 | 0.011 |
| rs3960984 | rs6804758 | 0.046 |
| rs3960984 | rs9822116 | 0.038 |
| rs3960984 | rs2111534 | 0.053 |
| rs4855093 | rs11719127 | 0.048 |
| rs4855093 | rs11920864 | 0.008 |
| rs4855093 | rs1976765 | 0.017 |
| rs4855093 | rs9814424 | 0.017 |
| rs4855093 | rs11706842 | 0.012 |
| rs4855093 | rs7641889 | 0.012 |
| rs4855093 | rs6786049 | 0.002 |
| rs4855093 | rs6799756 | 0.002 |
| rs4855093 | rs7645550 | 0.027 |
| rs3729679 | rs6443607 | 0.03 |
| rs3729679 | rs13092881 | 0.017 |
| rs3729679 | rs9879637 | 0.04 |
| rs3729679 | rs4501160 | 0.039 |
| rs3729679 | rs9841497 | 0.031 |
| rs3729679 | rs4583702 | 0.023 |
| rs3729679 | rs4955797 | 0.035 |
| rs3729679 | rs7652946 | 0.037 |
| rs3729679 | rs7612363 | 0.012 |
| rs3729679 | rs4579062 | 0.006 |
| rs3729679 | rs7639391 | 0.016 |
| rs3729679 | rs6790867 | 0.015 |
| rs3729679 | rs1542 | 0.017 |
| rs3729679 | rs7633318 | 0.013 |
| rs3729679 | rs6793893 | 0.016 |
| rs3729679 | rs6800015 | 0.028 |
| rs3729679 | rs4955807 | 0.021 |
| rs3729679 | rs6772028 | 0.032 |
| rs3729679 | rs6784495 | 0.044 |
| rs3729679 | rs2677770 | 0.046 |
| rs3729679 | rs12494623 | 0.001 |
| rs12488074 | rs9841497 | 0.015 |
| rs12488074 | rs4583702 | 0.017 |
| rs12488074 | rs4955797 | 0.014 |
| rs12488074 | rs6791364 | 0.019 |
| rs12488074 | rs7629064 | 0.036 |
| rs12488074 | rs6784495 | 0.018 |
| rs12488074 | rs9968179 | 0.022 |
| rs12488074 | rs2699887 | 0.027 |
| rs12488074 | rs2699905 | 0.23 |
| rs12488074 | rs2677770 | 0.017 |

**Supplementary Table 3:** Summary of METABRIC and TCGA datasets

| **Factor** | **METABRIC** | **TCGA** |
|---|---|---|
| Total Number | 94 | 178 |
| | *Median (LQ, UQ)* | *Median (LQ, UQ)* |
| **Age at diagnosis** (years) | 60.61 (50.06, 69.62) | 58.13 (49.00, 67.00) |
| **Follow-up all cases** (years) | 9.0 (4.2, 13.7) | 1.1(0.2, 2.9) |
| **Follow-up still living** (years) | 12.2 (7.9, 17.4) | 0.9 (0.2, 2.6) |
| **Vital status** | | |
| Alive | 47 (50%) | 153 (86%) |
| Dead | 47 (50%) | 25 (14%) |
| **Tumour size** | 22 (17.18, 28.75) | - |
| **Lymph nodes positive** | | |
| Number (0, 1, 2, >3) | 50, 11, 12, 21 | 69, 27, 15, 38 |
| **Grade** | | |
| I | 13 (13.8%) | - |
| II | 35 (37.2%) | - |
| III | 44 (46.8%) | - |
| **Stage** | | |
| I | 31 (33%) | 56(31%) |
| II | 27 (28.8%) | 97 (54%) |
| III | 8 (8.5%) | 17 (9.6%) |
| IV | 2 (2.1%) | 8 (4.5%) |
| Not reported | 26 (27.6%) | - |
| **PAM50 subtype** | | |
| Basal | 7 (7.4%) | 7 (3.9%) |
| HER2 | 14 (14.9%) | 16 (9%) |
| Luminal A | 40 (42.6%) | 109 (61.2%) |
| Luminal B | 25 (26.6%) | 39 (21.9%) |
| Normal | 8 (8.5%) | 5 (2.8%) |
| Not reported | | 2 (1.1%) |
| **iCluster** | | |
| iC1 | 2 (2.1%) | - |
| iC2 | 6 (6.4%) | - |
| iC3 | 22 (23.4%) | - |
| iC4 | 13 (13.8%) | - |
| iC5 | 6 (6.4%) | - |
| iC6 | 2 (2.1%) | - |
| iC7 | 12 (12.8) | - |
| iC8 | 16 (17.0%) | - |
| iC9 | 10 (10.6%) | - |
| iC10 | 5 (5.3%) | - |
| **DNAseq** | | |
| reference allele reads | 79.5 (51.75, 124.75) | 68 (41, 112.25) |
| alternativeallele reads | 31.5 (20, 50) | 31.5 (20, 57) |
| **RNAseq** | | |
| reference allele reads | 78 (44.25, 16.25) | 43 (29.25, 61.75) |
| alternativeallele reads | 39 (23.25, 56.75 | 24 (14.25, 42.75) |

**Supplementary Table 4:** Summary Statistic of PIK3CA's MADE Survival Analysis in TCGA set. Median, LCL, and UCL in days; LCL: lower confidence level; UCL: upper confidence level; N: number of people of each group. (ER = estrogen; PR = progesterone; Her2 = human epidermal growth factor 2 receptor)

| **Discovery** | Overall Survival | | | | | | | Disease Specific Survival | | | | |
|---|---|---|---|---|---|---|---|---|---|---|---|---|
| | N | Events | Median | 0.95LCL | 0.95UCL | | | N | Events | Median | 0.95LCL | 0.95UCL |
| **MADE vs. no_MADE** | | | | | | | **MADE vs. no_MADE** | | | | | |
| no_MADE | 46 | 21 | 6750 | 4138 | NA | | no_MADE | 46 | 9 | NA | NA | NA |
| MADE | 45 | 26 | 2907 | 2112 | NA | | MADE | 45 | 18 | NA | 2582 | NA |
| **MADE groups** | | | | | | | **MADE groups** | | | | | |
| MADE_low | 6 | 4 | 1382 | 1293 | NA | | MADE_low | 6 | 3 | 1382 | 1293 | NA |
| no_MADE | 46 | 21 | 6750 | 4138 | NA | | no_MADE | 46 | 9 | NA | NA | NA |
| MADE_high | 39 | 22 | 3617 | 2149 | NA | | MADE_high | 39 | 15 | NA | 2907 | NA |
| **ER positive** | | | | | | | **ER positive** | | | | | |
| MADE_low | 5 | 3 | 2737 | 1293 | NA | | MADE_low | 5 | 2 | NA | 1293 | NA |
| no_MADE | 42 | 18 | 6750 | 4341 | NA | | no_MADE | 42 | 8 | NA | NA | NA |
| MADE_high | 28 | 12 | 4550 | 3617 | NA | | MADE_high | 18 | 7 | NA | NA | NA |
| **ER neg** | | | | | | | **ER neg** | | | | | |
| MADE_low | 1 | 1 | 1382 | NA | NA | | MADE_low | 1 | 1 | 1382 | NA | NA |
| no_MADE | 4 | 3 | 2539 | 744 | NA | | no_MADE | 4 | 1 | NA | 744 | NA |
| MADE_high | 11 | 10 | 985 | 939 | NA | | MADE_high | 11 | 8 | 1374 | 939 | NA |
| **PR pos** | | | | | | | **PR pos** | | | | | |
| MADE_low | 5 | 3 | 2737 | 1293 | NA | | MADE_low | 5 | 2 | NA | 1293 | NA |
| no_MADE | 37 | 14 | 6750 | 4680 | NA | | no_MADE | 37 | 7 | NA | NA | NA |
| MADE_high | 22 | 10 | 4550 | 3617 | NA | | MADE_high | 22 | 6 | NA | 3617 | NA |
| **PR neg** | | | | | | | **PR neg** | | | | | |
| MADE_low | 1 | 1 | 1382 | NA | NA | | MADE_low | 1 | 1 | 1382 | NA | NA |
| no_MADE | 9 | 7 | 3660 | 1296 | NA | | no_MADE | 9 | 2 | NA | 1799 | NA |
| MADE_high | 17 | 12 | 1530 | 962 | NA | | MADE_high | 17 | 9 | 2149 | 985 | NA |
| **HER2 pos** | | | | | | | **HER2 pos** | | | | | |
| MADE_low | 0 | | | | | | MADE_low | 0 | 0 | | | |
| no_MADE | 6 | 3 | 4138 | 2048 | NA | | no_MADE | 6 | 2 | NA | 2048 | NA |
| MADE_high | 9 | 6 | 2149 | 1071 | NA | | MADE_high | 9 | 5 | 2149 | 1530 | NA |
| **HER2 neg** | | | | | | | **HER2 neg** | | | | | |
| MADE_low | 6 | 4 | 1382 | 1293 | NA | | MADE_low | 6 | 3 | 1382 | 1293 | NA |
| no_MADE | 40 | 18 | 6750 | 4277 | NA | | no_MADE | 40 | 7 | NA | NA | NA |
| MADE_high | 30 | 16 | 3950 | 2907 | NA | | MADE_high | 30 | 10 | NA | 3447 | NA |

**Supplementary Table 5:** Bivariate association analysis between mutant allelic expression imbalance ratios and clinical data. Kruskal-Wallis rank sum tests were performed, and p-values were corrected per study for multiple testing using the Bonferroni method. (iC – Integrative Clusters)

| x | y | statistic | p_value | p_value_adjusted | parameter | method |
|---|---|---|---|---|---|---|
| METABRIC | | | | | | |
| Grade | alpha | 5.8086698 | 0.0547852165 | 0.657422598 | 2 | Kruskal-Wallis rank sum test |
| Stage | alpha | 0.8661607 | 0.8335853568 | 1.000000000 | 3 | Kruskal-Wallis rank sum test |
| iC | alpha | 21.7109523 | 0.0098416052 | 0.118099262 | 9 | Kruskal-Wallis rank sum test |
| PAM50 | alpha | 12.5258906 | 0.0138404490 | 0.166085388 | 4 | Kruskal-Wallis rank sum test |
| Age | alpha | 91.4026859 | 0.4684250452 | 1.000000000 | 91 | Kruskal-Wallis rank sum test |
| Lymph Node Count | alpha | 12.4720029 | 0.4893705632 | 1.000000000 | 13 | Kruskal-Wallis rank sum test |
| Grade | gamma | 1.3573920 | 0.5072780399 | 1.000000000 | 2 | Kruskal-Wallis rank sum test |
| Stage | gamma | 4.7326697 | 0.1924521221 | 1.000000000 | 3 | Kruskal-Wallis rank sum test |
| iC | gamma | 12.3712759 | 0.1931784163 | 1.000000000 | 9 | Kruskal-Wallis rank sum test |
| PAM50 | gamma | 13.4081293 | 0.0094445581 | 0.113334697 | 4 | Kruskal-Wallis rank sum test |
| Age | gamma | 91.9410974 | 0.4526694103 | 1.000000000 | 91 | Kruskal-Wallis rank sum test |
| Lymph Node Count | gamma | 9.2729374 | 0.7520315236 | 1.000000000 | 13 | Kruskal-Wallis rank sum test |
| TCGA | | | | | | |
| Stage | alpha | 11.3301843 | 0.0786922387 | 0.629537909 | 6 | Kruskal-Wallis rank sum test |
| PAM50 | alpha | 21.8470722 | 0.0002149679 | 0.001719743 | 4 | Kruskal-Wallis rank sum test |
| Age | alpha | 59.0136070 | 0.2059390303 | 1.000000000 | 51 | Kruskal-Wallis rank sum test |
| Lymph Node Count | alpha | 18.8712313 | 0.4651244044 | 1.000000000 | 19 | Kruskal-Wallis rank sum test |
| Stage | gamma | 5.8172040 | 0.4439755966 | 1.000000000 | 6 | Kruskal-Wallis rank sum test |
| PAM50 | gamma | 4.5303071 | 0.3389689529 | 1.000000000 | 4 | Kruskal-Wallis rank sum test |
| Age | gamma | 56.1253927 | 0.2887781190 | 1.000000000 | 51 | Kruskal-Wallis rank sum test |
| Lymph Node Count | gamma | 28.8974098 | 0.0676160384 | 0.540928307 | 19 | Kruskal-Wallis rank sum test |
